# Timing of treatment shapes the path to androgen receptor signaling inhibitor resistance in prostate cancer

**DOI:** 10.1101/2024.03.18.585532

**Authors:** Eugine Lee, Zeda Zhang, Chi-Chao Chen, Danielle Choi, Aura C. Agudelo Rivera, Eliot Linton, Yu-jui Ho, Jillian Love, Justin LaClair, John Wongvipat, Charles L. Sawyers

**Affiliations:** Human Oncology and Pathogenesis Program, Memorial Sloan Kettering Cancer Center, New York, NY, USA; Cancer Biology and Genetics Program, Memorial Sloan Kettering Cancer Center, New York, NY, USA; Howard Hughes Medical Institute, Chevy Chase, Maryland, USA

## Abstract

There is optimism that cancer drug resistance can be addressed through appropriate combination therapy, but success requires understanding the growing complexity of resistance mechanisms, including the evolution and population dynamics of drug-sensitive and drug-resistant clones over time. Using DNA barcoding to trace individual prostate tumor cells *in vivo*, we find that the evolutionary path to acquired resistance to androgen receptor signaling inhibition (ARSI) is dependent on the timing of treatment. In established tumors, resistance occurs through polyclonal adaptation of drug-sensitive clones, despite the presence of rare subclones with known, pre-existing ARSI resistance. Conversely, in an experimental setting designed to mimic minimal residual disease, resistance occurs through outgrowth of pre-existing resistant clones and <u>not</u> by adaptation. Despite these different evolutionary paths, the underlying mechanisms responsible for resistance are shared across the two evolutionary paths. Furthermore, mixing experiments reveal that the evolutionary path to adaptive resistance requires cooperativity between subclones. Thus, despite the presence of pre-existing ARSI-resistant subclones, acquired resistance in established tumors occurs primarily through cooperative, polyclonal adaptation of drug-sensitive cells. This tumor ecosystem model of resistance has new implications for developing effective combination therapy.

## INTRODUCTION

Cancer patients treated with chemotherapy, radiotherapy, molecularly targeted therapy and immunotherapy often confront the formidable challenge of drug resistance (1). These resistant tumors compromise patient prognosis and pose a significant obstacle to effective cancer management. The term “upfront”” resistance is often used to refer to clinical scenarios where the tumor is unresponsive at the beginning of treatment, whereas “acquired” resistance refers to tumors that are initially responsive but subsequently relapse. For cancers with canonical oncogenic driver mutations, upfront resistance may be a consequence of co-occurring genomic alterations that mitigate dependence of tumor cells on the initial primary driver alteration. *PTEN* loss in the setting of treating *PIK3CA*-mutant breast cancer with a PI3K inhibitor serves as an example (2). On the other hand, acquired resistance is a consequence of a cell-intrinsic or cell-extrinsic change that prevents an initially effective drug from continuing to inhibit tumor growth. Examples of cell intrinsic mechanisms include mutation of the drug target in a manner that preserves oncogenic activity but precludes inhibition, or epigenetic changes that alter the cell state of the tumor such that it is no longer dependent on the drug target (3–6). Cell-extrinsic mechanisms include changes in the tumor microenvironment (TME) that “rescue” the tumor by providing new growth signals that bypass the effect of the targeted therapy (7). More complex, population-level processes may also be at play, including transcriptional heterogeneity among tumor cells as well as adaptive TME responses to therapeutic stress that, collectively, can rewire survival pathways in tumor cells (8, 9).

The androgen receptor (AR) is a lineage survival factor for luminal prostate epithelial cells and, on that basis, provides the rationale for the use of androgen receptor signaling inhibitors (ARSI) as the primary treatment for advanced prostate cancer (10). ARSI therapy is used to treat patients with metastatic disease measured by radiographic imaging, as well as patients with biochemical relapse, a state of minimal residual disease defined by elevated levels of serum prostate specific antigen (PSA) but no radiographically detectable disease. Current insight into acquired ARSI resistance comes primarily from genomic sequencing analysis of tumors from patients with metastatic castration resistant prostate cancer (mCRPC). Mechanisms include AR amplification/mutation, structural variants in the AR genomic locus, and aberrant AR mRNA slicing, all of which typically reflect persistent AR pathway dependency. Of note, an increasing fraction of ARSI-resistant tumors display features of lineage plasticity, whereby tumor cells transition from a luminal state to basal-like, mesenchymal, stem-like or neuroendocrine states that are no longer dependent on AR (4, 6, 11–13). Whether these same mechanisms are relevant in the setting of minimal residual disease, where the surrounding TME is not fully established, is not known.

To gain insight into this question, we used *in vivo* barcoding to trace the evolutionary path resistance in two human prostate cancer xenografts treated with ARSI in two different ways: one to model the minimal residual disease setting and the second to model the measurable disease setting (4, 6, 7, 14–17). Remarkably, the path to ARSI resistance differs across these two treatment settings but the underlying molecular mechanisms are convergent. Furthermore, acquired resistance in the measurable disease setting requires cooperativity between subclones, indicative of a tumor ecosystem that could potentially be targeted to achieve effective combination therapy.

## RESULTS

### ARSI resistance in established tumors is adaptive despite the presence of pre-existing resistant clones

To determine the evolutionary trajectory of ARSI resistance (ARSI^R^), we performed *in vivo* cellular barcoding experiments to track individual cells in mixed populations using ClonTracer, a method previously used to study the origin of drug-resistant clones in models of non-small lung cancer and chronic myeloid leukemia (18) (19, 20). This method provides a powerful tool to distinguish between pre-existing resistance (the same barcode is enriched in multiple independent ARSI^R^ tumors but not in control tumors) and adaptive resistance (a similar barcode profile is recovered from ARSI^R^ and control tumors). This distinction between pre-existing and adaptive resistance can be statistically quantified using Pearson correlation matrices. Barcoding also provides an opportunity, through single cell cloning, to isolate the clones responsible for subsequent *in vivo* biology from the initial starting population of cells, thereby providing an opportunity to characterize the molecular properties responsible for emergent ARSI^R^ prior to exposure to any selective pressure.

We focused initially on the LNCaP/AR xenograft model due to its track record of revealing clinically relevant mechanisms of antiandrogen resistance (17). To introduce the library into LNCaP/AR cells, we intentionally used a low multiplicity of infection (MOI) to generate ∼10,000 uniquely barcoded cells, a number chosen based on an estimated frequency of pre-existing resistant cells (if they exist) of ∼0.1% and the need to recover identical barcodes from such pre-existing resistant cells across different xenografted animals (**Supplementary Fig. 1A**, see methods for more detail) (21–23). Toward this end, we generated a population of LNCaP/AR cells labeled with 11,600 unique barcodes (hereafter called pre-graft), expanded the cells, and then established xenografts in hormonally intact mice (control) or surgically pre-castrated mice treated with Enz (called pre-engraftment ARSI) to mirror the clinical use of ARSI therapy in a setting of non-radiographically measurable, minimal residual disease (**Figs. 1A-B**).

**Figure 1.**
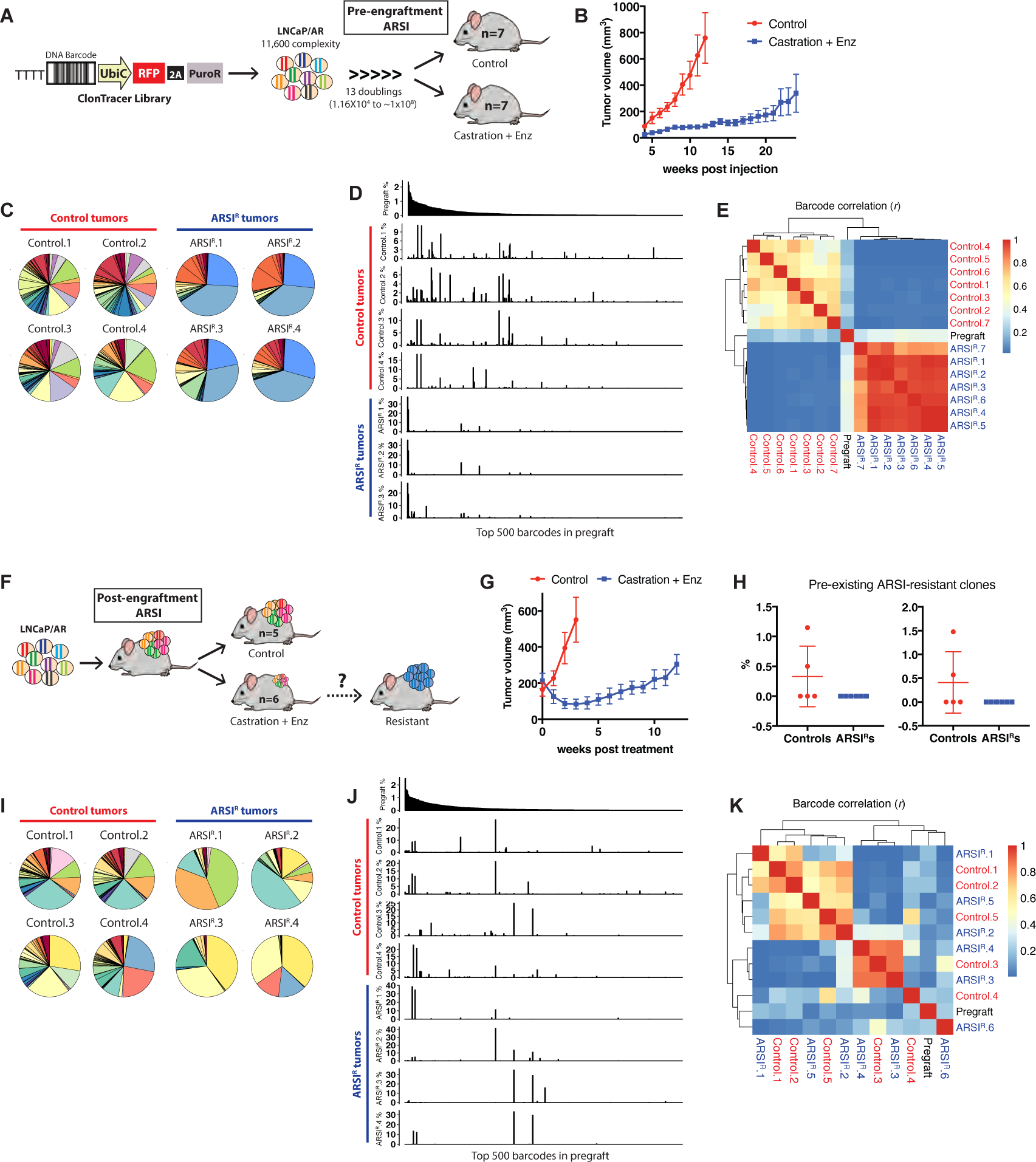
Evolutionary paths to ARSI resistance in LNCaP/AR xenograft model. (A) The LNCaP/AR barcode cell line was generated by lentiviral infection with ClonTracer library and grafted to hormonally intact mice (control) or physically pre-castrated mice treated with enzalutamide (Enz) (pre-engraftment ARSI). (B) Mean tumor volumes ± SEM (N=8 in each group). (C-D) Pie charts (C) and bar graphs (D) showing barcode distribution in the control and ARSI-resistant (ARSI^R^) tumors. In bar graphs, each bar represents each barcode with y-axis showing relative ratio, and the x-axes are identical across all the graphs in a decreasing order of abundance of 500 most enriched barcodes in pregraft (cell population used for xenograft, top graph). Barcode distribution of the full tumor set can be found in Supplementary Fig. 1. (E) Heatmap depicting Pearson correlation analysis (r) of barcodes between the tumors and hierarchical clustering on the values. (F) The LNCaP/AR barcode cell line was grafted to hormonally intact mice and when the tumors reached ∼200mm^3^, half of the mice were physically castrated and treated with Enz (post-engraftment ARSI) to test if the pre-existing ARSI-resistant clones will be selected in ARSI^R^ tumors. (G) Mean tumor volumes ± SEM after Enz treatment (N=6 in control and N=7 in Enz treated group). (H) The relative ratio of two pre-existing ARSI-resistant clones enriched in control and ARSI^R^ tumors. (I-J) Pie charts (I) and bar graphs (J) showing barcode distribution in control and ARSI^R^ tumors. Barcode distribution of the full tumor set can be found in Supplementary Fig. 3. (K) Heatmap depicting Pearson correlation analysis (r) of barcodes between the tumors and hierarchical clustering on the values.

After tumor progression, control and ARSI-resistant (ARSI^R^) tumors were harvested and subjected to next-generation sequencing (NGS) to compare barcode recovery across these two conditions relative to pre-graft. As expected, the number of barcodes recovered from tumors in both groups was reduced relative to pre-graft, reflecting a well-known bottleneck in tumor initiation seen when xenografting cancer cell lines. Turning to the comparison of interest, ARSI^R^ tumors contained significantly fewer barcodes than controls, and these barcodes were shared across tumors (**Fig. 1C, Supplementary Figs. 1B-C**). To quantify the difference in barcode enrichment in the intact versus ARSI setting, we ranked each barcode by percentage abundance and noted that barcode enrichment in control versus ARSI^R^ tumors was non-overlapping (**Fig. 1D and Supplementary Fig. 1D**). This conclusion was confirmed by a Pearson correlation analysis showing distinct control and ARSI^R^ clusters (**Fig. 1E**). Thus, ARSI resistance in the pre-engraftment setting is caused by outgrowth of pre-existing resistant subclones.

The two dominant clones enriched in all ARSI^R^ tumors (average 33.4% and 23.6%, blue pie slices in ARSI^R^ tumors, **Fig. 1C and Supplementary Fig. 1C**) were detected at 2.1% and 1.7% respectively in the pre-graft (blue bars in top graph in **Supplementary Fig. 1E**), and thus were enriched 10-20 fold under the selective pressure of ARSI treatment. Conversely, these clones were 4-5 fold depleted in control tumors (average 0.57% and 0.35%), suggesting their growth may be inhibited in hormonally intact mouse. We explored this question by adding dihydrotestosterone (DHT) or Enz to our standard pre-graft culture media for 7 days and examined changes in the global barcode distribution (**Supplementary Figs. 2A-B**). The abundance of the two dominant ARSI^R^ tumors clones in the pre-graft (blue bars in **Supplementary Fig. 1E**) was decreased by DHT treatment (**Supplementary Fig. 2C**), indicative of androgen-dependent growth suppression.

The pre-engraftment ARSI treatment scenario described above was selected to model the clinical scenario of ARSI treatment when patients have minimal residual disease, e.g., those with a rise in serum PSA level (called biochemical relapse or BCR). To model the use of ARSI in patients with measurable disease, we injected the barcoded LNCaP/AR cells into hormonally intact mice, waited until tumors reached ∼200mm^3^, then administered ARSI therapy (castration + Enz) (hereafter called post-engraftment ARSI) **(Fig. 1F).** As expected from earlier work using this model (4), tumors grown in this setting responded initially to ARSI but then progressed, at which time we compared barcode distribution between control and ARSI^R^ tumors (**Fig. 1G**). As with the pre-ARSI experiment (**Supplementary Fig. 1B**), the number of enriched barcodes in ARSI^R^ tumors was lower than control tumors (**Supplementary Fig. 3A**); however, the resistant clones enriched in the pre-engraftment ARSI setting (reflecting minimal residual disease) were surprisingly <u>depleted</u> in the post-engraftment ARSI setting (measurable disease) (average 0.00076% and 0.00082%, **Fig. 1H**). Rather, we found a common set of barcodes enriched in both control and ARSI^R^ tumors, indicating that clones in control tumors adapted to ARSI treatment and were sustained or further enriched (**Figs. 1I-J, Supplementary Figs. 3B-C**). The barcode correlation analysis showed that control and ARSI^R^ tumors were no longer separated (**Fig. 1K**), revealing a distinct evolutionary path from that seen in the pre-engraftment ARSI setting (**Fig. 1E**). Instead, they were grouped into three main clusters, each of which contain both ARSI^R^ <u>and</u> control tumors. Furthermore, >90% of the barcodes in the post-engraftment ARSI experiment were also enriched in control tumors (yellow data points in scatter plots and bar graphs in **Supplementary Fig. 3D**). Thus, the clones that gave rise to control tumors persisted and adapted to ARSI treatment. Surprisingly, the small number of pre-existing, drug resistant subclones that cause resistance the pre-engraftment setting did not expand.

To determine if this finding extends to an independent model, we performed an analogous barcode labeling xenograft experiment using the CWR22Pc cell line (**Fig. 2A and Supplementary Fig. 4A**, see details in methods) (7). Unlike the LNCaP/AR model, CWR22Pc cells did not progress to ARSI resistance despite >5 months of therapy. Remarkably however, tumors arose within 2-3 weeks of discontinuing Enz treatment, indicating that tumor cells in this model persist at the injection site for months and are poised to resume growth if the selective pressure of ARSI is partially mitigated by Enz removal (**Fig. 2B and Supplementary Fig. 5A**). Cognizant of this difference between CWR22Pc and LNCaP/AR in the pre-engraftment ARSI setting (persistence versus resistance), we nonetheless analyzed the relative distribution of barcodes in control versus ARSI^R^ tumors exactly as before. As we saw with LNCaP/AR, a small number of clones present in the pre-graft were highly enriched in ARSI^R^ tumors, indicative of a pre-existing resistance mechanism. The only difference was modest enrichment of three of these barcodes in control tumors, likely indicative of the high engraftment potential of these clones even in the absence of ARSI treatment. Importantly, the fact that control and ARSI^R^ tumors clustered separately in barcode correlation analysis underscores that resistance in the pre-engraftment ARSI setting is pre-existing (**Figs. 2C-E, Supplementary Figs. 4B-D, 5B-D**).

**Figure 2.**
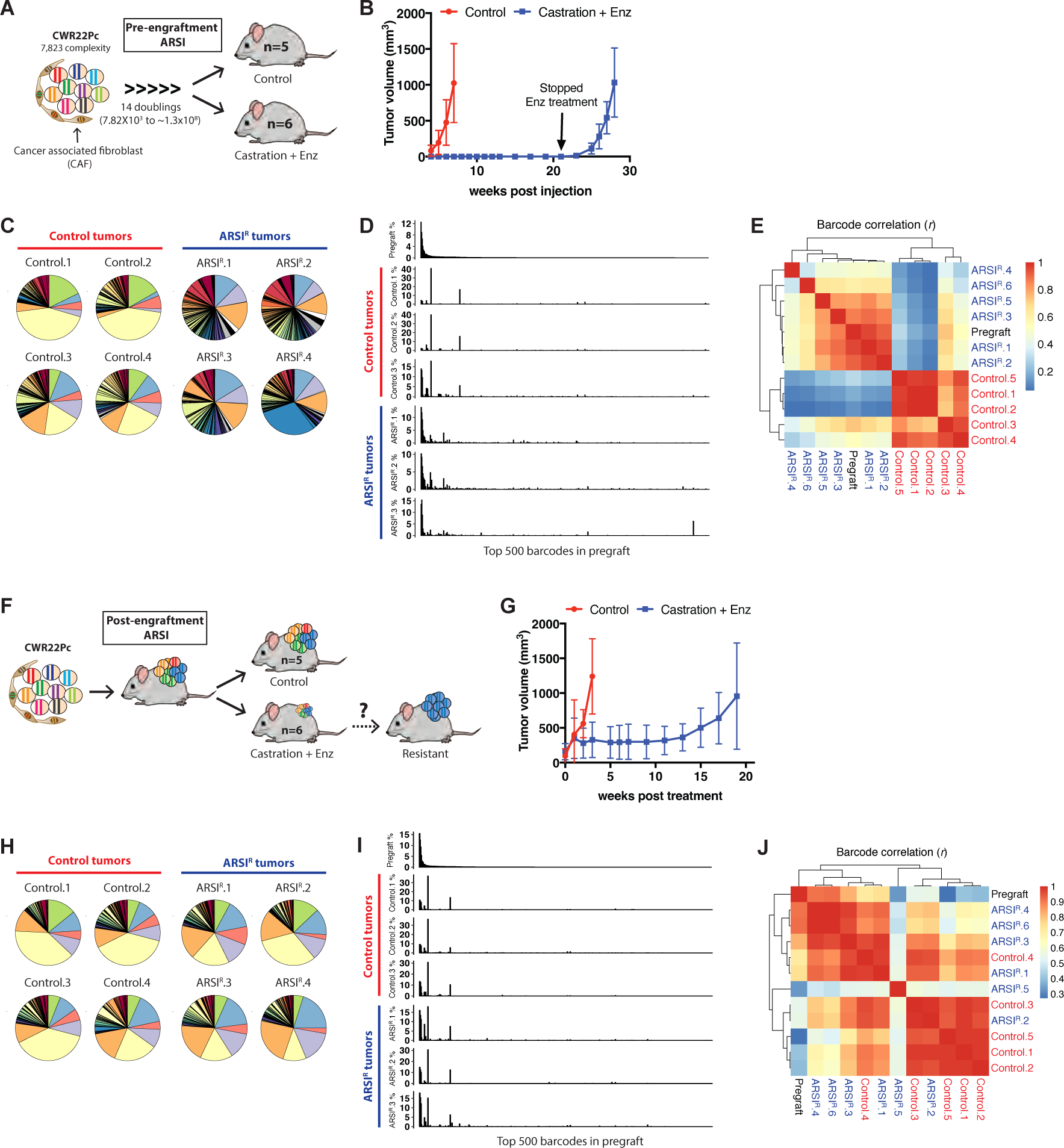
Evolutionary paths to ARSI resistance in CWR22Pc xenograft model. (A) The CWR22Pc cells containing cancer associated fibroblast (CAF) were infected with lentivirus containing ClonTracer library and grafted to hormonally intact mice (control) or physically pre-castrated mice treated with enzalutamide (Enz) (pre-engraftment ARSI). (B) Mean tumor volumes ± SEM (N=10 in control and N=8 in Enz treated group). Note that the Enz treatment is stopped at 21 weeks after grafting. (C-D) Pie charts (C) and bar graphs (D) showing barcode distribution in control and ARSI-resistant (ARSI^R^) tumors. In bar graphs, the barcodes in x-axes are identical across all the graphs in a decreasing order of abundance of 500 most enriched barcodes in pregraft. Barcode distribution of the full tumor set can be found in Supplementary Fig. 7. (E) Pearson correlation analysis (*r*) of barcodes between the tumors shows that the control and ARSI^R^ tumors are clustered separately. (F) The CWR22Pc barcode cell line was grafted into hormonally intact mice and when the tumors reached ∼200mm^3^, half of the mice were physically castrated and treated with Enz (post-engraftment ARSI). (G) Mean tumor volumes ± SEM after Enz treatment (N=5 in control and N=6 in Enz treated group). (H-I) Pie charts (H) and bar graphs (I) showing barcode distribution in control and ARSI^R^ tumors. Barcode distribution of the full tumor set can be found in Supplementary Fig. 6. (J) Pearson correlation analysis (*r*) of barcodes between the tumors shows that the control and ARSI^R^ tumors are clustered together.

Having documented pre-existing resistance in two models when ARSI therapy is given prior to tumor engraftment, we next asked if a similar switch to adaptive resistance occurs in the CWR22Pc model when treatment is started after tumor engraftment (**Fig. 2F**). As expected, established CWR22Pc tumors responded to ARSI treatment by entering a cytostatic phase then developed acquired resistance within 10-20 weeks (**Fig. 2G and Supplementary Fig. 7A**). Barcode distribution graphs and correlation analysis showed intermixing of barcodes in control and ARSI^R^ tumors in the established tumor setting, in contrast to the pre-engraftment ARSI setting where the barcode distribution was separated (**Figs. 2H-J and Supplementary Figs. 6A-C, 7B-D**). Furthermore, the percentage of shared barcodes between control and ARSI^R^ tumors was higher when ARSI was given post-engraftment compared to pre-engraftment (Venn diagrams in **Supplementary Figs. 6D and 7E**) and the relative abundance of these shared barcodes is ∼90% in both groups in the post-engraftment setting (yellow data points in scatter plots and bar graphs in **Supplementary Figs. 6D and 7E**). In summary, barcode sequencing analysis of two different prostate cancer xenograft models shows that the path to ARSI resistance differs depending on the timing of ARSI therapy. In a minimal residual disease setting (modeled by pre-engraftment ARSI treatment), a small number of subclones contribute to resistance. However, in a measurable disease setting (modelled by post-engraftment ARSI treatment), resistance primarily evolves though adaptation of the dominant clones present in ARSI-sensitive tumors.

The sharing of barcodes in control and ARSI^R^ tumors provides conclusive evidence that acquired resistance in the established tumor setting is adaptive, a mechanism generally viewed as a consequence of epigenetic rewiring or perturbation of feedback circuits (24, 25). To determine if genetic mechanisms may also play a role, we performed targeted exome sequencing of DNA from ARSI^R^ tumors using a comprehensive 468 gene panel (MSK IMPACT) (26). As expected, baseline and acquired genomic alterations were seen in both models, but none were shared across resistant tumors with the exception of a single missense mutation in PTPRD (D767R) detected in 4 of 4 ARSI^R^ CWR22Pc tumors, but at a low allele burden (average variant allele frequency = 0.09) (**Supplementary tables 1-2**). Although PTPRD has been reported as a potential tumor suppressor (27), the D767R mutation has not been reported in existing cancer genome databases (cBioPortal). Thus, while the barcoding data provides clear evidence that acquired resistance in the established tumor setting is adaptive, further work is needed to conclusively eliminate any genetic contribution.

### Pre-existing and adaptive paths to ARSI resistance converge on common mechanisms

Having demonstrated these different paths to resistance through lineage tracing, we next turned our attention to the underlying molecular causes within each path. In the post-engraftment setting, we previously reported glucocorticoid receptor (GR) signaling and HER3 kinase activation (by stromal production of NRG1) as the primary molecular drivers of ARSI resistance in LNCaP/AR and CWR22PC, respectively (4, 7). Furthermore, we found that GR expression (in LNCaP/AR cells) and NRG1 expression (in stromal cells) are both upregulated by ARSI therapy *in vivo*, consistent with the adaptive response confirmed by the current barcoding experiments. Now armed with the barcode identity of the specific clones responsible for ARSI resistance *in vivo*, we sought to isolate these clones from the mixture of cells in the pre-graft as this would allow us to characterize them before exposure to ARSI, i.e., before any selective pressure that might drive an adaptive response.

Through single cell cloning and barcode sequencing of several hundred clonal isolates from the LNCaP/AR pre-graft, we identified two clones that contributed substantially to *in vivo* ARSI resistance in the pre-engraftment setting (hereafter called pre-ARSI^R1^ and pre-ARSI^R2^). We also isolated one clone enriched in control tumors (pre-Intact) and four neutral clones that were not enriched in any tumors (clones A-D) (**Fig. 3A**). To determine whether the two pre-ARSI^R^ clones had any evidence of pre-existing ARSI resistance, we characterized their *in vitro* growth characteristics in the presence or absence of Enz. Pre-ARSI^R1^ and pre-ARSI^R2^ both displayed resistance to Enz (2-3 fold increase in IC50) compared to the neutral (A-D) and pre-Intact clones (**Fig. 3B and Supplementary Figs. 8A-B**). In addition, both pre-ARSI^R^ clones had elevated baseline AR transcriptional output measured by a commonly used AR reporter (**Supplementary Figs. 8C-D**).

**Figure 3.**
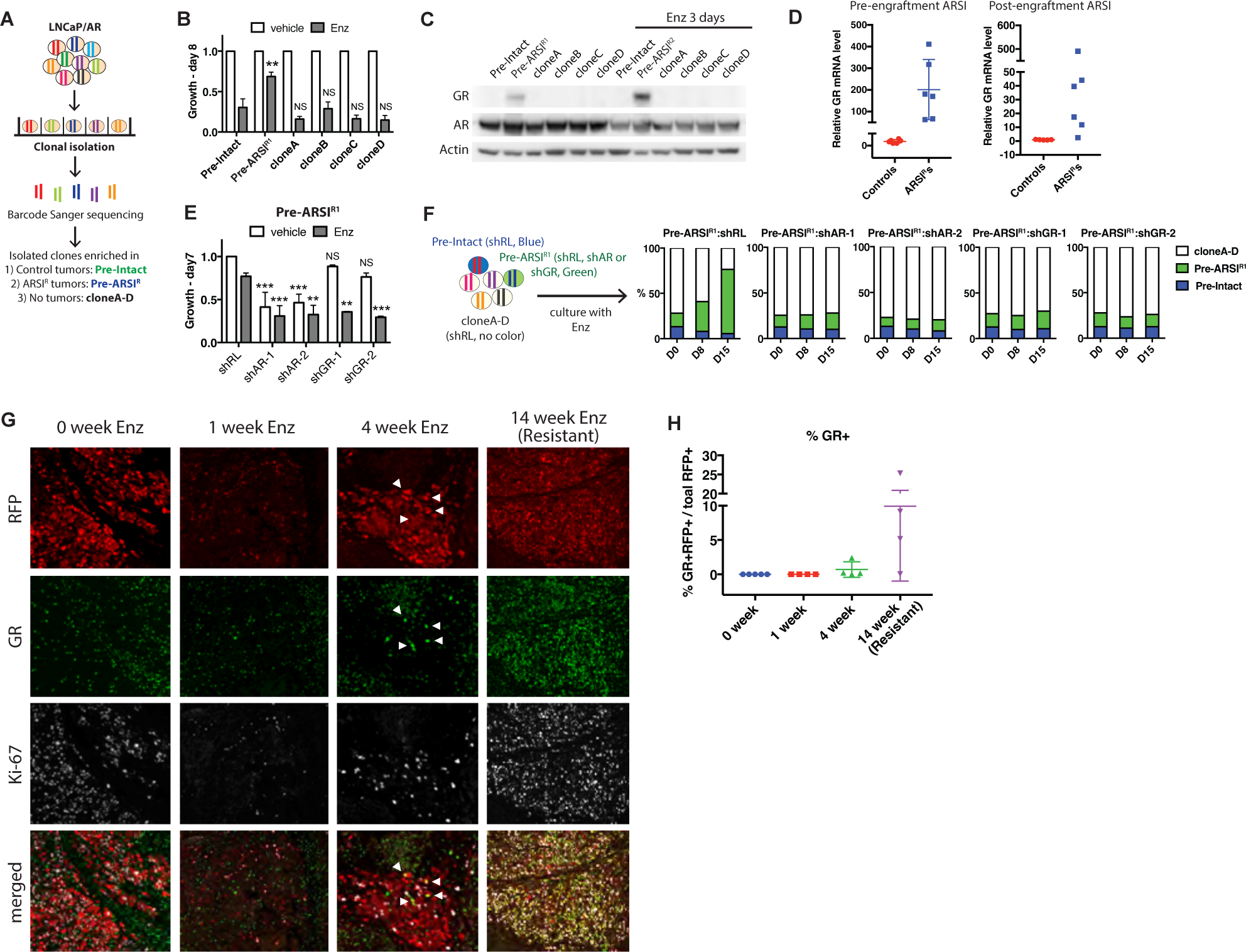
Evolutionary paths to ARSI resistance in LNCaP/AR model converge on GR upregulation. (A) The clonal isolation was performed by sorting single cell of LNCaP/AR barcode cell line into 96-well plate and sanger sequencing of barcode in each clonal line. One clone enriched across control tumors (Pre-Intact), one clone enriched across ARSI^R^ tumors in pre-engraftment ARSI setting (Pre-ARSI^R^), and four clones not enriched in the tumors (cloneA through D) were isolated. (B) The clones were cultured in regular media with vehicle or hormone-deprived media with 10μM Enz for 8 days. The cell viability values of Enz treated cells were normalized to the values of vehicle treated cells ± SD of two biological replicates. *P*-values were determined by one-way ANOVA compared to the value of Enz treated Pre-Intact clone. (C) GR is upregulated in Pre-ARSI^R1^ clone and 3 days of 10μM Enz treatment further increased GR level in Pre-ARSI^R1^ clone. (D) qRT-PCR shows that GR mRNA levels (mean ± SD) are upregulated in ARSI^R^ tumors derived from pre-engraftment (Fig. 1B) and post-engraftment (Fig. 1G) ARSI studies. (E) Knockdown of AR or GR re-sensitizes Pre-ARSI^R1^ clone to 10μM Enz. The cell viability values were normalized to the value of vehicle treated shRenilla (shRL) ± SD of two biological replicates. *P*-values were determined by two-way ANOVA compared to shRL. (F) The GFP-labeled Pre-ARSI^R1^ clone was infected with shRL, shAR or shGR and mixed with Azurite-labeled Pre-Intact clone and clonesA-D infected with shRL. The mixed population was then cultured with 10μM Enz and the relative ratio of cells with blue, green and no color were analyzed using flow cytometry at day 0, 8 and 15. (G) The LNCaP/AR barcode cell line was grafted into hormonally intact mice and when the tumors reached ∼400mm^3^, the mice were physically castrated and treated with Enz. The tumors were collected at 0, 1 and 4 weeks after treatment, and at 14 weeks when became resistant to the treatment, and stained with RFP (red, grafted LNCaP/AR cells), GR (green) and Ki-67 (grey). (H) The numbers of RFP+/GR+ cells were normalized to the numbers of total RFP+ cells in each tissue section. Mean ± SD, N=4 in each group.

Having previously implicated GR as the driver of adaptive resistance *in vivo* (4), we asked if GR might also be responsible for pre-existing resistance. We first confirmed GR is highly expressed in both pre-engraftment ARSI^R^ and post-engraftment ARSI^R^ tumors *in vivo* (**Fig. 3D**). We then performed RNA sequencing of the *in vitro* pre-graft clones and found that GR is 1.8-fold upregulated in the pre-ARSI^R1^ clone compared to pre-Intact and clones A-D (**Supplementary table 3**). Furthermore, both pre-ARSI^R^ clones express GR protein that is further induced by just 3 days of Enz treatment (**Fig. 3C and Supplementary Fig. 8E**), a finding reminiscent of the “GR-primed” state reported previously in cell lines isolated from LNCaP/AR xenografts with acquired ARSI resistance (28). To determine if baseline or Enz-induced GR expression in the pre-ARSI^R^ clones is the cause of Enz resistance, we examined the consequences of GR and AR shRNA knockdown on cell growth. GR knockdown restored Enz sensitivity to the pre-ARSI^R^ clones, while AR knockdown blocked cell growth regardless of Enz treatment. (**Fig. 3E, Supplementary Figs. 8F-H**). To mimic the emergence of drug resistance in a polyclonal population, we mixed fluorescently labeled pre-ARSI^R^ clones (green) expressing control shRNA (Renilla, RL), AR shRNA or GR shRNA, the pre-Intact clone (blue) and the 4 neutral A-D clones (no color) and performed an *in vitro* cell growth competition assay in Enz. AR and GR knockdown, but not the RL control, blocked expansion of the Pre-Enz clone (**Fig. 3F**), providing further evidence that GR upregulation is the driver of pre-existing ARSI resistance in this model.

With this insight documenting baseline GR expression in rare pre-ARSI^R^ clones prior to engraftment, we re-examined GR expression in the established LNCaP/AR tumor model, but now in a time course experiment (0, 1, 4, 14 weeks) to monitor the kinetics of GR induction following initiation of ARSI therapy. Because the barcode analysis in this setting revealed that resistance does not emerge from pre-ARSI^R^ clones (**Fig. 1F**), we were interested in the kinetics of GR expression across the entire population of tumor cells, particularly at early time points when one would expect rare pre-existing resistant clones to expand. To ensure precision in this assessment, we took advantage of RFP expression in the engrafted LNCaP/AR cells to avoid scoring GR expression in mouse endothelial and stromal cells. Specifically, we counted GR+/RFP+ (double positive) cells at each time point, together with Ki67 to track proliferation. As expected, the number of Ki-67+ cells was low (but clearly measurable) 1 week after ARSI treatment, increased at 4 weeks and fully evident at 14 weeks (**Fig. 3G**). Notably, despite quantitative immunofluorescence analysis of tens of thousands of cells, no GR+/RFP+ tumor cells were detectable after 1 week but were clearly evident at 4 and 14 weeks (**Figs. 3G-H**). We conclude that pre-ARSI^R^ clones, which are present (albeit rare) in intact tumors, fail to expand after ARSI therapy. However, adaption of pre-Intact clones is evident within 4 weeks.

Having documented a shared mechanism (GR upregulation) in the different paths to resistance in the LNCaP/AR model, we asked if the same theme applies to the CWR22Pc model, where NRG1/HER3 is the primary mechanism of adaptive resistance *in vivo* (7). As with LNCaP/AR, we isolated and screened single cell clones from the CWR22Pc pre-graft and identified the highest enriched clone responsible for Enz persistence (pre-ARSI^P^) as well as three neutral clones (clones A-C) (**Fig. 4A**). We previously reported that *in vitro* growth of CWR22Pc tumor cells is responsive to conditioned media (CM) derived from cancer associated fibroblasts (7). We therefore evaluated the *in vitro* growth of the pre-ARSI^P^ clone and the three neutral clones in media with or without CM, as well as in presence or absence of DHT. All four clones grew at similar rates in the absence of CM; however, the pre-ARSI^P^ clone expanded more quickly in CM in a dose-dependent manner, in regular and hormone-deprived media (**Figs. 4B-C**). Consistent with earlier work documenting HER3 activation by NRG1 as the mechanism of CM action in these cells (7), CM treatment led to dose-dependent phosphorylation of HER3 and AKT in the pre-ARSI^P^ clone but not in the neutral clones (**Fig. 4D**). Furthermore, treatment with the dual HER2/HER3 inhibitor neratinib reversed the growth promoting effect of CM in the pre-ARSI^P^ clone but had no effect on the neutral clones (29) (**Fig. 4E**). *In vivo* phospho-HER3 and phospho-AKT levels were also higher in ARSI^R^ tumors compared to controls, as well as NRG1 expression (**Figs. 4F-G**). Taken together, the evidence from both models of ARSI resistance demonstrates that the molecular basis of resistance in each model (GR in LNCaP/AR; NRG1/HER3 in CWR22Pc) is conserved despite the different evolutionary paths that tumor cells undertake in the pre-engraftment versus post-engraftment setting.

**Figure 4.**
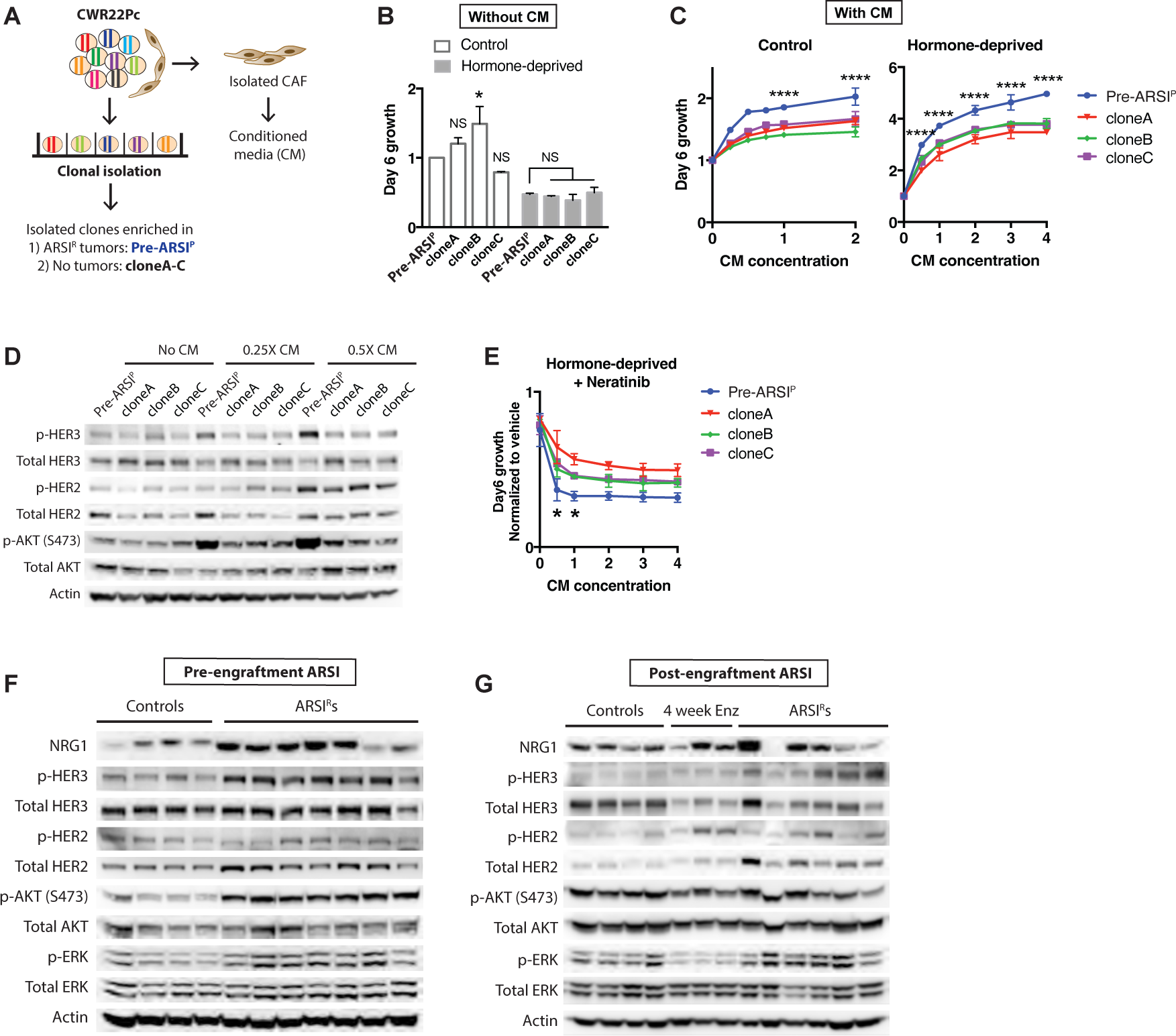
Evolutionary paths to ARSI resistance in CWR22Pc model converge on HER3. (A) One clone enriched across the ARSI^R^ tumors in pre-engraftment ARSI setting (Pre-ARSI^P^), three clones not enriched in the tumors (cloneA through C) and CAFs were isolated from CWR22Pc barcode cell line. (B) The clones were cultured in control or hormone-deprived media for 6 days. The cell viability values in control media were normalized to the value of Pre-ARSI^P^ ± SD and the values in hormone-deprived media were normalized to the values in control media ± SD of two biological replicates. NS = not significant, \**p*<0.05, two-way ANOVA compared to Pre-ARSI^P^. (C) The clones were cultured in regular or hormone-deprived media with increasing concentration of conditioned media (CM) derived from CAF for 6 days. The cell viability values were normalized to the value of no CM ± SD of two biological replicates. \*\*\*\**p*<0.0001, two-way ANOVA compared to cloneA. (D) The clones were cultured in serum-free media for 2hrs and cultured with each indicated concentration of conditioned media (CM) for 10min. (E) The clones were cultured in hormone-deprived media with 0.01μM neratinib and increasing concentration of conditioned media (CM) derived from CAF for 6 days. The cell viability values were normalized to the value of no CM ± SD of two biological replicates. (F) NRG1 and total HER2, and phospho-HER3, AKT and ERK levels are increased in ARSI^R^ tumors from pre-engraftment ARSI study (Fig. 2B). (G) NRG1 and total HER2, and phospho-HER2, HER3 and ERK levels are increased in some of the ARSI^R^ tumors from post-engraftment ARSI study (Fig. 2G).

### The adaptive enzalutamide resistance requires inter-clonal cooperation

To gain further insight into how established ARSI-sensitive tumors adapt to ARSI treatment, we repeated the LNCaP/AR *in vivo* barcoding experiment described in Figure 1 but now using a greatly simplified pool consisting only of the 6 single cell clones derived from the LNCaP/AR pre-graft (**Figs. 5A-B**). Given our earlier evidence that the relative fraction of pre-existing ARSI^R^ cells in the bulk pre-graft is reduced by DHT (**Supplementary Fig. 2B**), we intentionally increased the percentage of the pre-ARSI^R^ clone in the pool by 3-fold to further enhance the potential for this clone to contribute to ARSI resistance *in vivo*. As expected from the original pre-graft experiment using the entire pool (**Fig. 1C**), the pre-Intact clone was highly enriched in control tumors and the pre-ARSI^R^ clone was highly enriched in the pre-engraftment ARSI setting (**Fig. 5A**). However, in the post-engraftment ARSI setting, the pre-Intact clone was massively enriched in both control <u>and</u> ARSI^R^ tumors despite our attempt to “stack the odds” for the pre-ARSI^R^ clone in the pre-graft (**Fig. 5B**). Furthermore, the ARSI^R^ tumors derived from the pre-Intact clone which do not express GR in vitro now express high levels of GR *in vivo* (**Fig. 5C**), documenting their remarkable capacity to undergo adaptive resistance.

**Figure 5.**
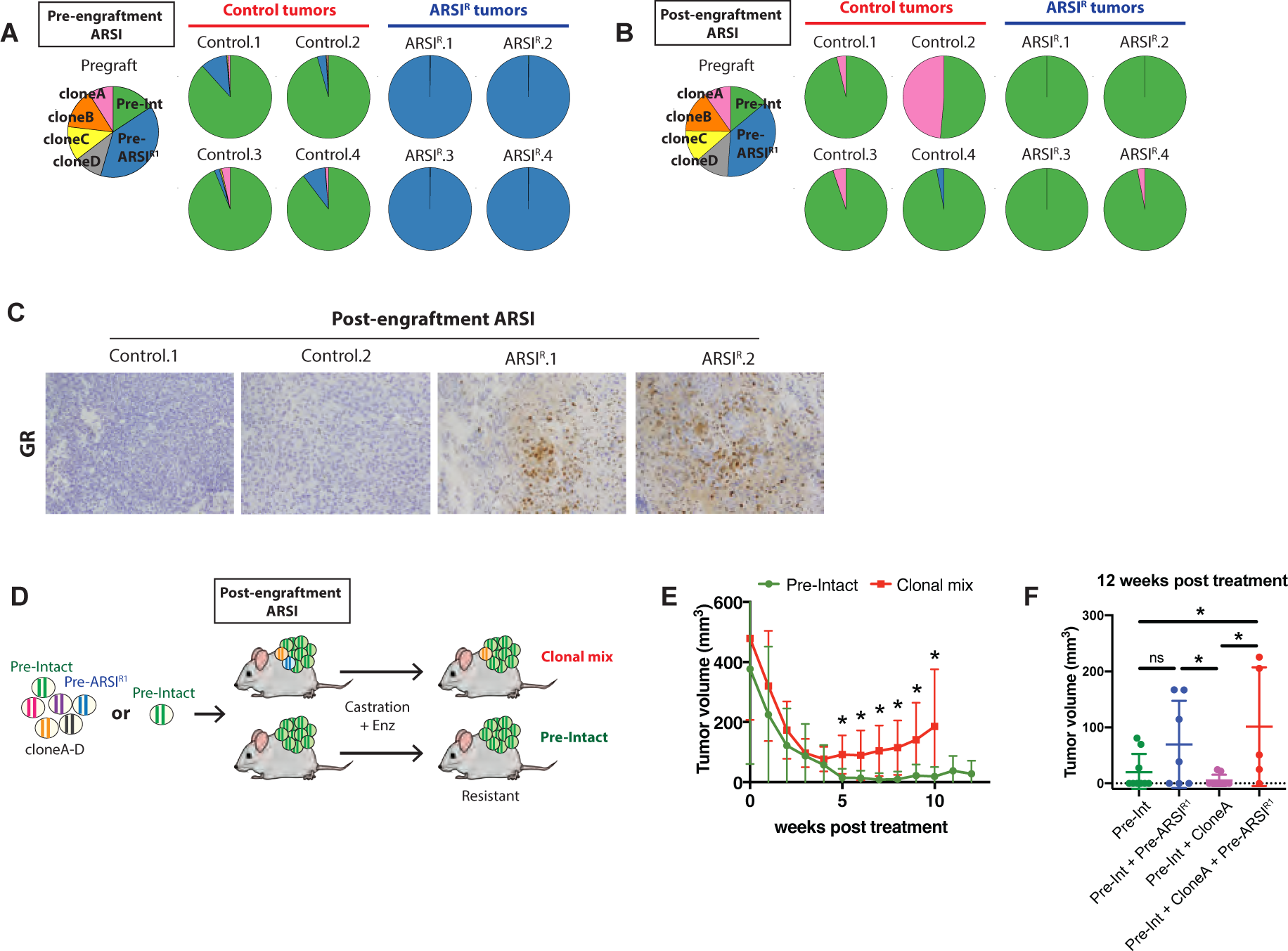
ARSI resistance in established tumors requires inter-clonal cooperation. (A-B) Mix of 6 LNCaP/AR clones (Pre-Intact, Pre-ARSI^R1^ and clonesA-D) were grafted to control, pre-engraftment ARSI treated (A), or post-engraftment ARSI treated (B) mice. Barcode distribution in pregraft, control and ARSI^R^ tumors are shown in pie charts. (C) GR protein levels were accessed in control and ARSI^R^ tumors in B. (D) Mix of 6 LNCaP/AR clones or Pre-Intact clone alone were grafted to post-engraftment ARSI setting. (E) Mean tumor volumes ± SEM after ARSI treatment (N=7 in each group). *P*-value was determined by two-tailed *t*-test at each time point. (F) Mix of indicated clones were grafted to post-engraftment ARSI setting. Tumor volumes were measured 12 weeks after ARSI treatment (N=9 in Pre-Intact and Pre-Intact + CloneA, N=7 in Pre-Intact + Pre-ARSI^R1^, and N=5 in Pre-Intact + CloneA + Pre-ARSI^R1^ group). Mean tumor volumes ± SEM. *P*-value was determined by two-tailed *t*-test. For all panels, NS = not significant, \**p*<0.05.

To further investigate this capacity, we simplified the adaptive resistance model even further by repeating the experiment with just the pre-Intact clone alone. As expected, pre-Intact cells efficiently gave rise to tumors in control mice; however, they were no longer capable of adaptive resistance when injected alone (**Figs. 5D-E, Supplementary Fig. 9A**). In light of recent evidence of cooperativity between subclones in promoting tumor progression, metastasis and drug resistance in other models (5, 30–32), we asked if clonal cooperativity might be contributing to adaptive ARSI resistance. We paired the pre-Intact clone with either the pre-ARSI^R^ clone, clone A (selected based on its capacity to form small tumors in intact mice when injected alone (**Supplementary Fig. 9A**)) or both. Remarkably, the ability of pre-Intact cells to undergo adaptive ARSI resistance was now restored but only when paired with the pre-ARSI^R^ clone (with or without clone A) (**Fig. 5F**). Thus, clonal interaction is required for adaptive resistance in this model. Whether this cooperativity occurs through paracrine mechanisms or physical cell-to-cell interaction will require further study.

## DISCUSSION

Acquired resistance to targeted cancer therapy is a dynamic process that evolves with treatment through a mix of genetic and non-genetic mechanisms (33–36). The recent application of high-content lineage tracing has revealed the impact of transcriptional heterogeneity in facilitating this evolution (32, 37–40). Here we have applied this technology to prostate cancer, using *in vivo* barcoding to track the evolutionary path to ARSI resistance, but adding a new variable of modeling resistance by varying the timing of ARSI treatment. When ARSI therapy was administered early (prior to tumor engraftment, to mirror the clinical scenario minimal residual disease), only a few clones were capable of resistance and, in all cases, that resistance was pre-existing. However, when ARSI was initiated after tumors had already established, drug-sensitive tumor cells were the primary contributors to resistance - despite the fact that the same pre-existing resistant clones that cause resistance in the minimal residual disease setting are present in the established tumors. Furthermore, the adaptation of these drug sensitive clones to ARSI resistance required cooperativity with other tumor clones that, in the end, represented a minor fraction of the emergent drug resistant tumor.

To our knowledge, timing of therapy has not been directly evaluated as a variable that influences the path to treatment resistance. The fact that established tumors develop resistance through an adaptive path is likely a consequence of increased heterogeneity (due to larger tumor burden) as well as the surrounding ecosystem of stromal and inflammatory cells that are not present to the same degree in the setting of minimal residual disease. Importantly, the experimental model reported here provides a tractable system to dissect the molecular details of the adaptive response. For example, the barcoding analysis reveals that ∼35% of drug-sensitive clones in an established tumor consistently contribute to resistance. This suggests that not all ARSI-sensitive cells in untreated tumors are capable of adaptation to ARSI therapy. By comparing the “adaptable” and “non-adaptable” populations from the pre-graft populations of each model, it should be possible to gain insight into the molecular basis of adaptability. In the LNCaP/AR model reported here, adaptability is likely explained by a unique chromatin state in “adaptable” cells that is permissive for GR upregulation in response to ARSI therapy, as suggested by an earlier study documenting AR repression of a GR enhancer in GR-positive LNCaP/AR subline (28). A similar mechanism could explain enhanced sensitivity to stromal NRG1 in a subset of CWR22Pc tumor cells, perhaps through elevated expression HER kinase receptors.

Another critical question is how cooperativity among clones in established tumors promotes adaptability. A similar phenomenon has been described in breast cancer, where cytokine-dependent communication of “neutral” tumor clones with infiltrating myeloid cells helps drive the growth of the resistant tumor clone (41). In the prostate model described here, we have isolated “cooperativity” to a single “helper” clone which, coincidently, is the same clone responsible for pre-existing resistance in the minimal residual disease model. Curiously, despite their capacity to thrive in the setting of ARSI therapy, these inherently ARSI-resistant cells switch to a new role in tumors in hormonally intact mice, “helping” the more abundant population of ARSI-sensitive clones adapt to ARSI treatment. Having now reduced the complexity of clonal cooperativity to just two isolated tumor cell clones, it should be possible to dissect the molecular details of cooperativity and “help” through detailed a single cell analysis of the adaptive process over time. The answer could provide insight into strategies to disrupt the “helper” cell signal and thereby prevent the emergence of adaptive resistance.

## Materials and Methods

### Cell lines

LNCaP/AR cell line was generated and maintained as previously described (14). CWR22Pc was a gift from Marja T. Nevalainen (Thomas Jefferson University, Philadelphia, PA) and maintained as previously described (1). All cells were tested negative for mycoplasma (Lonza, LT07-318).

### Barcode library transduction

The ClonTracer library was a kind gift from Novartis. Lentiviral transduction of cells was performed as previously described (1). 100,000 LNCaP/AR or CWR22Pc cells were plated in 6 well plate (Corning, 353046) and the attached number of cells in the extra well was counted on the following day before the infection. Cells were then infected with ClonTracer library at low multiplicity of infection (MOI) to enable each cell has one copy of reporter construct. Cells with stable integration of the library were measured by mCherry flow cytometry and selected with 1 μg/ml puromycin (Invivogen, ant-pr). The selected cells were expended in culture for 13-14 doublings to obtain enough number of cells for xenograft experiments.

### Xenograft assay

For all experiments, 2 × 10^6^ LNCaP/AR or CWR22Pc barcode cells were injected subcutaneously into the flank of CB17 SCID mice in a 50:50 mix of matrigel (Corning, 356237) and regular culture medium. The same number of cells used for injection (2 × 10^6^) was also snap-frozen at each xenograft experiment and referred to as ‘pre-graft’. Since the mouse-derived cancer associated fibroblast (CAF) in CWR22Pc are also labelled with barcode, we FACS sorted the human EpCAM-positive/mouse MHC-class1-negative population from the injected cells at each xenograft experiment and referred to it as ‘pre-graft’. Once tumors were palpable, tumor size was measured weekly using Peira TM900 system (Peira bvba, Belgium). For pre-engraftment androgen receptor signaling inhibitor (ARSI) experiments, cells were injected into hormonally intact or physically castrated mice 1 week prior to the injection, and enzalutamide treatment (10 mg/kg) was initiated on the day of injection to the castrated mice, 5 days a week by oral gavage. For post-engraftment ARSI experiments, cells were injected into the intact mice first and when the xenografts reached 200∼400mm^3^, mice were randomized and the half of the mice were physically castrated and treated with enzalutamide. Once these ‘resistant’ tumors exceeded their original volume, the mice were sacrificed, tumors were collected into tissue grinder (Fisher Scientific, 02-542-09), and the ground tissues were snap-frozen for analysis. All animal experiments were performed in compliance with the approved institutional animal care and use committee (IACUC) protocols of the Research Animal Resource Center of Memorial Sloan Kettering Cancer Center.

### Barcode sequencing and analysis

Genomic DNA was extracted from the frozen tissues with PureGene Core Kit A (Qiagen). Barcode sequences were PCR amplified for NGS by introducing Illumina adaptors and 10-bp-long index sequences. For each PCR reaction, 2 μg of genomic DNA was used as a template using Titanium Taq PCR kit (Takara). The sequence information for the primers used for barcode amplification was kindly provided by Novartis and can be found in Supplementary Table 4. The PCR-amplified products were quantified using PicoGreen, analyzed by Agilent TapeStation, and then run on a HiSeq 2500 in Rapid Mode in a 125bp/125bp paired end run, using the HiSeq Rapid SBS Kit v2 (Illumina). Libraries were also run on a NextSeq 500 in a 150bp/150bp paired end run, using the TG NextSeq 500/550 Mid Output Kit v2 (300 Cycles) (Illumina).

To analyze barcode composition, paired-end reads were joined together using FLASH tool (2). The joined reads were processed with FASTX-toolkit (3). Reads containing the constant region in the vector “ACTGACTGCAGTCTGAGTCTGACAG” were filtered. Then the adjacent 30 mer variable sequence were cropped and counted for frequencies. All the cropped sequences were filtered for the designed WS pattern. The sequences with frequency < 2 were further filtered out for downstream analysis.

### IMPACT sequencing

Genomic DNA extracted from the frozen tissues or cell pellets underwent genomic profiling with a hybridization capture–based next-generation sequencing assay (Integrated Mutation Profiling of Actionable Cancer Targets [MSK-IMPACT] 468) as previously described (4).

### Clonal isolation

Single clones were isolated from LNCaP/AR or EpCAM-positive/mouse MHC-class1-negative CWR22Pc barcode cells into 96-well plate using FACS. Barcode sequence of each clone was PCR amplified directly from the cells using Terra PCR Direct Polymerase kit (Takara) for Sanger sequencing using previously described primers (5).

### shRNA knockdown of AR and GR

For shRNA knockdown experiments, LT3GEPIR vector (all-in-one Tet-ON system, gift from Johannes Zuber) (6) with the following guide sequences were used:

shAR-1: TAGTGCAATCATTTCTGCTGGC

shAR-2: TTGAAGAAGACCTTGCAGCTTC

shGR-1: TCCAAAGCAGTTTCACTCTCAA

shGR-2: GAAGCTGTAAAGTTTTCTTCAA

shRenilla (shRL) was previously described as Ren.713 targeting Renilla luciferase (6). Cells with stable integration of the hairpin were selected with 1 μg/ml puromycin and also sorted by GFP flow cytometry after 3 days of 100 ng/ml doxycycline (Sigma) treatment.

### AR luciferase reporter assay

1 × 10^4^ cells/well were plated in triplicate on white 96 well plates (Corning, 3903) and cotransfected with SV40 Renilla Luciferase and ARR3-Luciferase constructs using Effectene Transfection Reagent (Qiagen). Luciferase activity was assayed 48 hr after transfection using Dual-Luciferase Reporter Assay System (Promega). To measure enzalutamide IC50, the cells were plated and transfected in hormone-deprived media (10% charcoal-stripped dextran-treated fetal bovine serum, Omega Scientific, FB-04). Cells were treated with DMSO or enzalutamide (0.1, 1 or 10 μM) on the following day, and luciferase activity was assayed 24 hr after treatment. IC50 values were determined using GraphPad Prism (GraphPad).

### Conditioned Media Collection

To isolate CAFs, 1 × 10^6^ CWR22Pc cells were trypsinized and re-suspended in 200 μl staining buffer (PBS containing 3% BSA). Cells were blocked for 30 min and stained with 2μl of human EpCAM (FITC conjugated, Miltenyi Biotec, 130-080-301) and 1:75 of mouse MHC class I (APC conjugated, BioLegend, 114613) antibodies at room temperature in the dark for 30 min. Cells were then washed 3 times, counterstained with DAPI (Invitrogen) and the FITC-negative/APC-positive population was sorted out using BD FACSAria cell sorter (BD Biosciences).

To collect conditioned media (CM), 4 × 10^6^ CAFs were plated in 10cm dish. The following day the media was removed and replaced with serum free media after washing the cells with PBS twice. CM was collected 3 days later, filtered with a 0.45μM filter (Millex, SLHA033SS), concentrated 10x using Vivaspin™ protein concentrator spin columns (Sartorius, VS15T02) and stored at 4°C.

### Cell growth assay

For all experiments, 1-2 × 10^3^ cells/well were plated in triplicate on white 96 well plates (Corning, 3903). Cells were culture in regular media or hormone-deprived media with indicated concentration of enzalutamide, DHT or DMSO for each experiment. CellTitier-Glo assay (Promega) was performed according to manufacturer’s instructions at indicated days and the viability was normalized to day 0 or DMSO. For clones with AR or GR knockdown, the cells were pre-treated with doxycycline for 3 days and plated for growth assay with continuous doxycycline treatment. To test the effect of CM on growth, CWR22Pc clones were plated in 100 μl regular or hormone-deprived media per well, and 100 μl CM with concentration twice higher than indicated was added to each well the next day.

### FACS-based competition assay

Pre-Intact clone was infected with pLV-Azurite (blue fluorescent protein, Addgene, 36086) and Pre-ARSI^R1^ clone was infected with SGEP-Renilla-713 (GFP) (4). Cells with stable integration of the fluorescent plasmids were sorted by flow cytometry. Equal number of Pre-Intact (blue), Pre-ARSI^R1^ (green) and cloneA-D (no color) were mixed and cultured with DMSO, 10 μM enzalutamide or 10 nM DHT. The relative ratio of blue, green and no color cells were analyzed using flow cytometry at indicated days.

To test the effect of AR or GR knockdown on the growth of Pre-ARSI^R1^ in mixed population, the GFP-labeled Pre-ARSI^R1^ was infected with shRL, shAR or shGR, and the Azurite-labeled Pre-Int and cloneA-D were infected with shRL. Cells with stable integration of the hairpin were selected with 1 μg/ml puromycin and equal number of Pre-ARSI^R1^ (green; shRL, shAR or shGR), Pre-Intact (blue; shRL) and cloneA-D (no color; shRL) were mixed and cultured with 10 μM enzalutamide. The relative ratio of blue, green and no color cells were assayed at indicated days using LSRII (BD Biosciences) and analysis was done using FlowJo software (Tree Star).

### Western Blot

Protein was extracted from cells or frozen tissues as previously described (7). Protein lysates were subjected to SDS-PAGE and immunoblotted with the following antibodies against: AR (Abcam, ab108341), GR (BD Transduction Laboratories, 611227), actin (Cell Signaling Technology, 4970), tubulin (Santa Cruz Biotechnology, sc-9104), Cyclophilin B (Abcam, ab178397), NRG1 (Cell Signaling, 2573), HER2 (Cell Signaling, 2165), pHER2 (Cell Signaling, 2243), HER3 (Cell Signaling, 12708), pHER3 (Cell Signaling, 4791), pAKT (Cell Signaling, 4060), AKT (Cell Signaling, 4691), pERK (Cell Signaling, 4370), or ERK (Cell Signaling, 9102).

### Immunostaining

The immunofluorescent staining was performed at MSKCC Molecular Cytology Core Facility using Discovery XT processor (Ventana Medical Systems). The tissue sections were deparaffinized with EZPrep buffer (Ventana Medical Systems), antigen retrieval was performed with CC1 buffer (Ventana Medical Systems). Sections were blocked for 30 minutes with Background Buster solution (Innovex), followed by avidin-biotin blocking for 8 minutes (Ventana Medical Systems). Multiplex immunofluorescent stainings were performed as previously described (8) with GR (Cell Signaling, 12041, 0.5μg/ml), Ki-67 (Abcam, ab15580, 1μg/ml) or tRFP (Evrogen, AB233, 1μg/ml) primary antibodies and Tyramide Alexa Fluor 488 (Invitrogen, B40953), Tyramide CF543 (Biotum, 92172) or Tyramide CF594 (Biotum, 92174) secondary antibodies, respectively. After staining, slides were counterstained with DAPI (Sigma, D9542, 5 μg/ml) for 10 min and coverslipped with Mowiol, and image was scanned. The number of total cells (DAPI+), RFP+ and RFP+/GR+ cells in each slide were counted using ImageJ. The average of total cells counted per slide was 86,000 cells.

The Immunohistochemistry staining was performed at MSKCC Pathology Core Facility. Tumor pieces were fixed in 4% PFA prior to paraffin embedding and then stained with GR (1:1000, Cell Signaling, D6H2L) on a bond III automated stainer (Leica Biosystems, IL).

### Transcription analysis

RNA was extracted from cells or frozen tissues as previously described (7). For quantitative PCR with reverse transcription (RT–qPCR), we used the High-Capacity cDNA Reverse Transcription Kit (Applied Biosystems) to synthesize cDNA according to the manufacturer’s protocol. Real-time PCR was performed using 2X SYBR green quantfast PCR Mix (Qiagen, 1044154) and gene-specific primers as follows: human-specific Actin and GR. Data were analyzed by the DDCT method using Actin as a control gene and normalized to control samples, which were arbitrarily set to 1. Primers for GR and ACTB were purchased through Qiagen. For RNA-seq, library preparation, sequencing and expression analysis were performed by the New York Genome Center. Libraries were prepared using TruSeq Stranded mRNA Library Preparation Kit in accordance with the manufacturer’s instructions and sequenced on an Illumina HiSeq2500 sequencer (rapid run v2 chemistry) with 50 base pair (bp) reads. To analyze RNA-seq data, reads were aligned to the NCBI GRCh37 human reference using STAR aligner (9). Quantification of genes annotated in Gencode vM2 were performed using featureCounts and quantification of transcripts using Kalisto (10). QC were collected with Picard and RSeQC (11) (http://broadinstitute.github.io/picard/). Normalization of feature counts was done using the DESeq2 package (http://www-huber.embl.de/users/anders/DESeq/).

### Statistics

Information on replication, statistical tests and presentation are given in the respective figure legends. For comparison of pooled data between two different groups, unpaired t tests were used to determine significance. For comparison of data among three groups, one-way or two-way ANOVA was used to determine significance. In all figures, *P<0.05, **P<0.01, ***P<0.001 and ****P<0.0001. Statistical significance analyses were performed using GraphPad Prism version 7.0.

## Acknowledgments

We thank the Integrated Genomics Operation Core at MSKCC, funded by the NCI Cancer Center Support Grant (CCSG, P30 CA08748), Cycle for Survival, and the Marie-Josée and Henry R. Kravis Center for Molecular Oncology, for conducting NGS and MSK-IMPACT. We thank MSKCC Molecular Cytology Core, funded by P30 CA008748, for assistance with IF staining. We also thank New York Genome Center for conducting the RNA-sequencing and analysis, MSKCC Pathology Core for assistance with IHC staining, MSKCC Flow Cytometry Core for technical support, Novartis for generously providing ClonTracer library and primer sequences, and the members of the Sawyers laboratory for helpful discussions.

---

**Supplementary figure 1.**
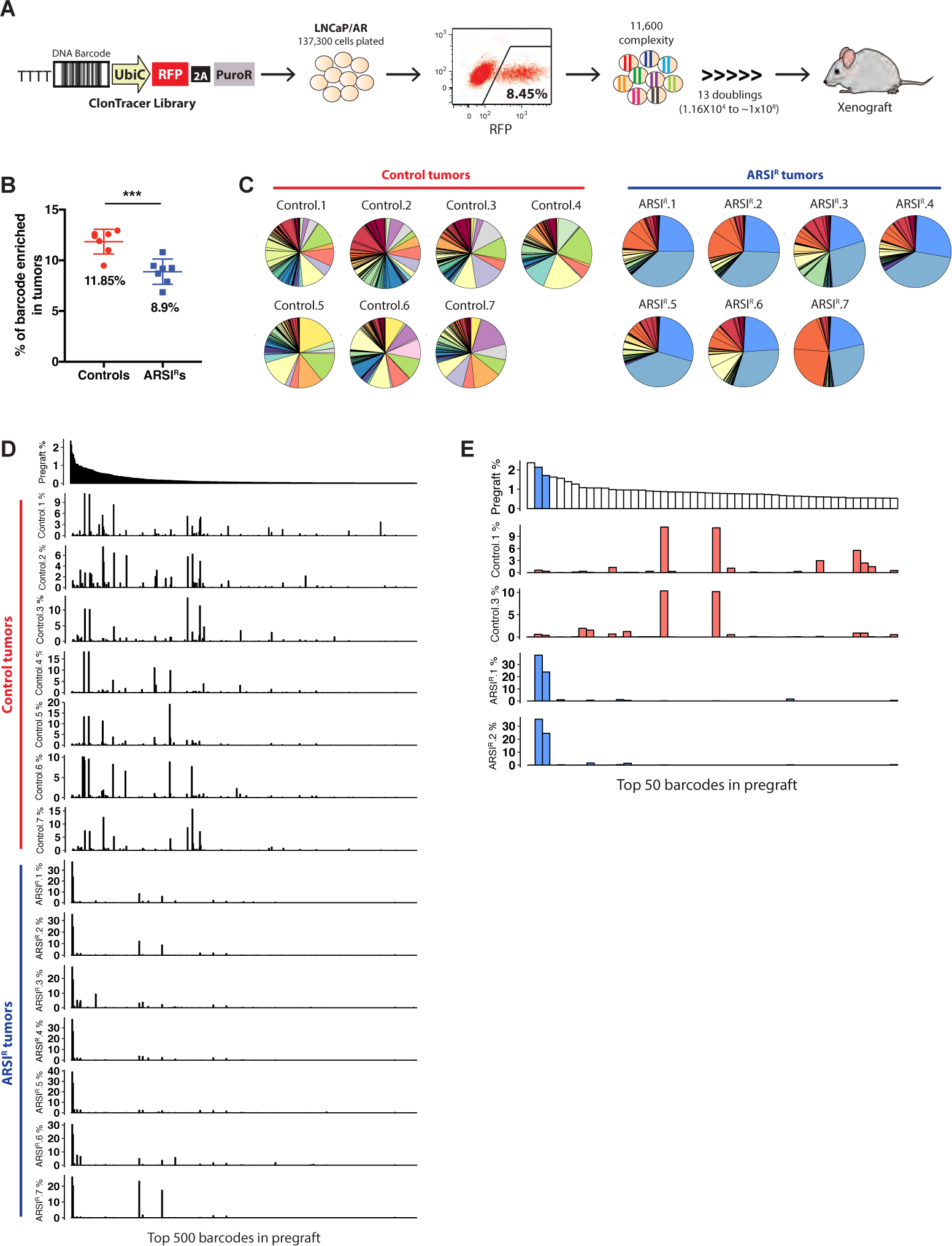
Enrichment of pre-existing resistance clones in pre-engraftment ARSI setting. (A) The LNCaP/AR barcode cell line was generated by lentiviral infection with ClonTracer library to enable 11,600 cells labeled with unique barcode sequence (details can be found in Materials and methods). The infected cells then went through 13 doublings and used for xenograft experiments. (B) Significantly lower number of barcodes are enriched in ARSI-resistant (ARSI^R^) tumors from Fig. 1B. Mean ± SD, N=7 in each group. \*\*\**p*<0.001, two-tailed *t*-test. (C-D) Pie charts (C) and bar graphs (D) showing barcode distribution in control and ARSI^R^ tumors. In bar graphs, each bar represents each barcode with y-axis showing relative ratio, and the x-axes are identical across all the graphs in a decreasing order of abundance of 500 most enriched barcodes in pregraft (cell population used for xenograft, top graph). (E) Distribution of the top 50 barcodes in pregraft shows that the barcodes most enriched in ARSI^R^ tumors (blue bars) are some of the most enriched barcodes in pregraft.

**Supplementary figure 2.**
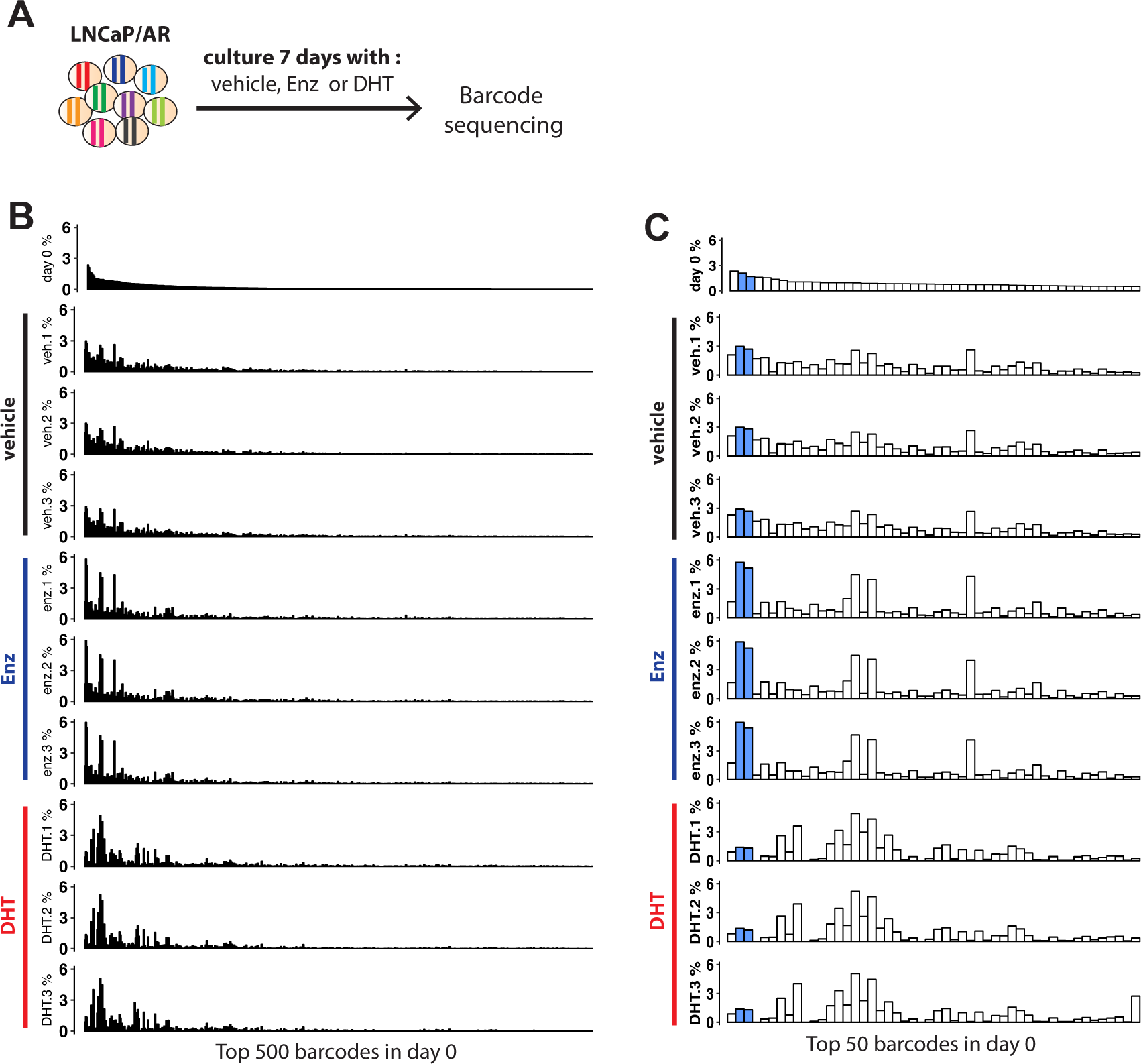
In vitro effects of AR activation and inhibition on growth of pre-existing ARSI-resistant clones. (A) LNCaP/AR barcode line was culture with either DMSO vehicle, 10μM Enz or 10nM DHT for 7 days, and then gDNA was extracted for barcode sequencing. (B-C) Distribution of the top 500 (B) or the top 50 (C) barcodes enriched in LNCaP/AR barcode cell line before the treatment (day 0, top graph) and in triplicates of each treatment group. The relative abundance of pre-existing ARSI-resistant clones (blue bars in Supplementary Fig. 1E and here) are increased by vehicle or Enz, and decreased by DHT treatment.

**Supplementary figure 3.**
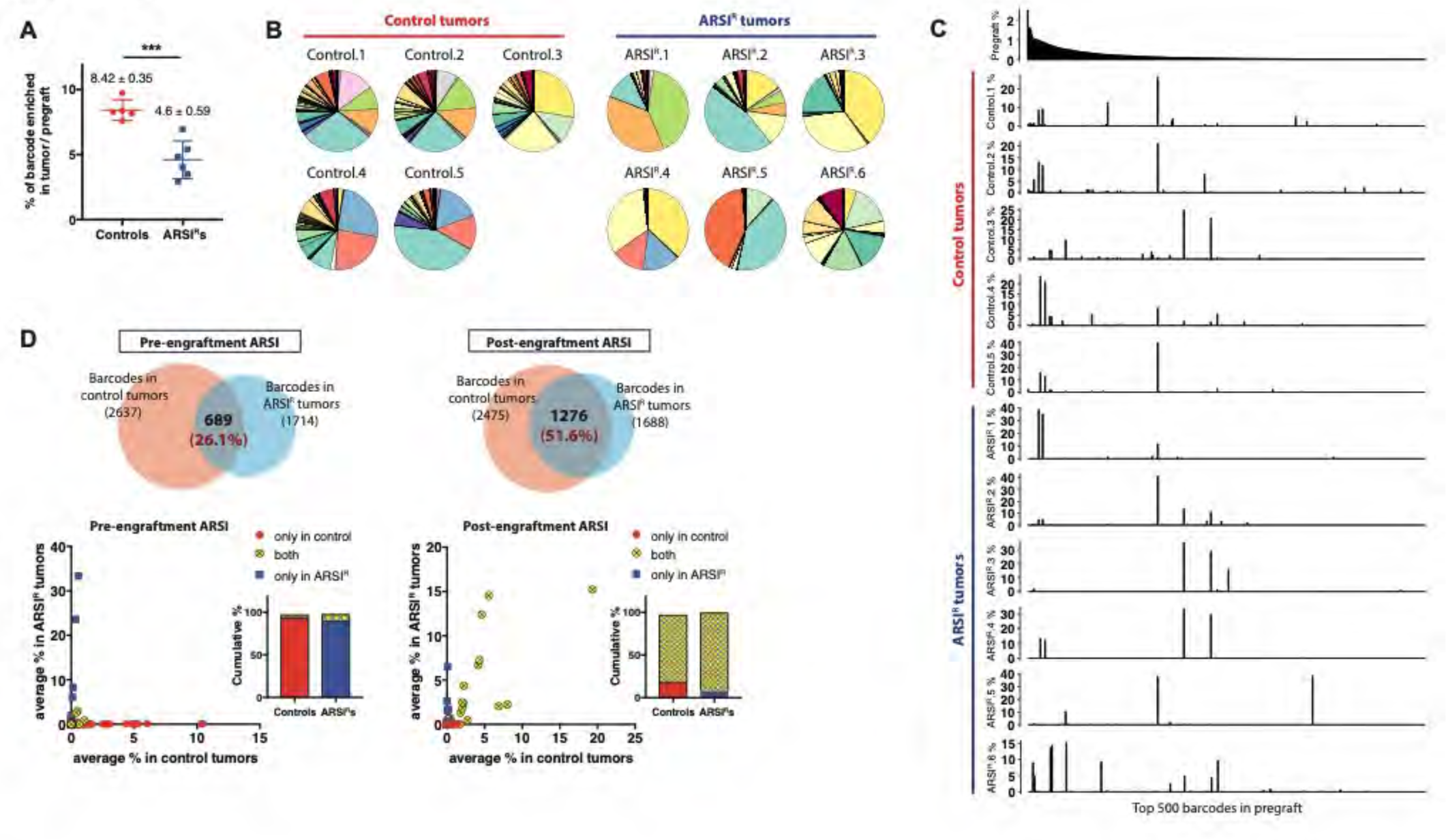
ARSI resistance in established LNCaP/AR tumors is adaptive despite the presence of pre-existing resistant clones. (A) Significantly lower number of barcodes are enriched in ARSI^R^ tumors from Fig 1G. N=5 in control and N=6 in ARSI^R^ group. \*\*\**p*<0.001, two-tailed *t*-test. (B-C) Pie charts (B) and bar graphs (C) showing barcode distribution in control and ARSI^R^ tumors. In bar graphs, the barcodes in x-axes are identical across all the graphs in a decreasing order of abundance of 500 most enriched barcodes in pregraft. (D) Top, Venn diagrams showing the overlaps between barcodes enriched in control and ARSI^R^ tumors in pre-engraftment ARSI (Fig. 1B) and post-engraftment ARSI (Fig. 1G) studies. The % overlap is based on the number of barcodes enriched in control tumors. Bottom, scatter plots showing the average % of each barcode in tumors and bar graphs showing cumulative % of barcodes enriched only in control (blue), ARSI^R^ (red) or both groups (yellow).

**Supplementary figure 4.**
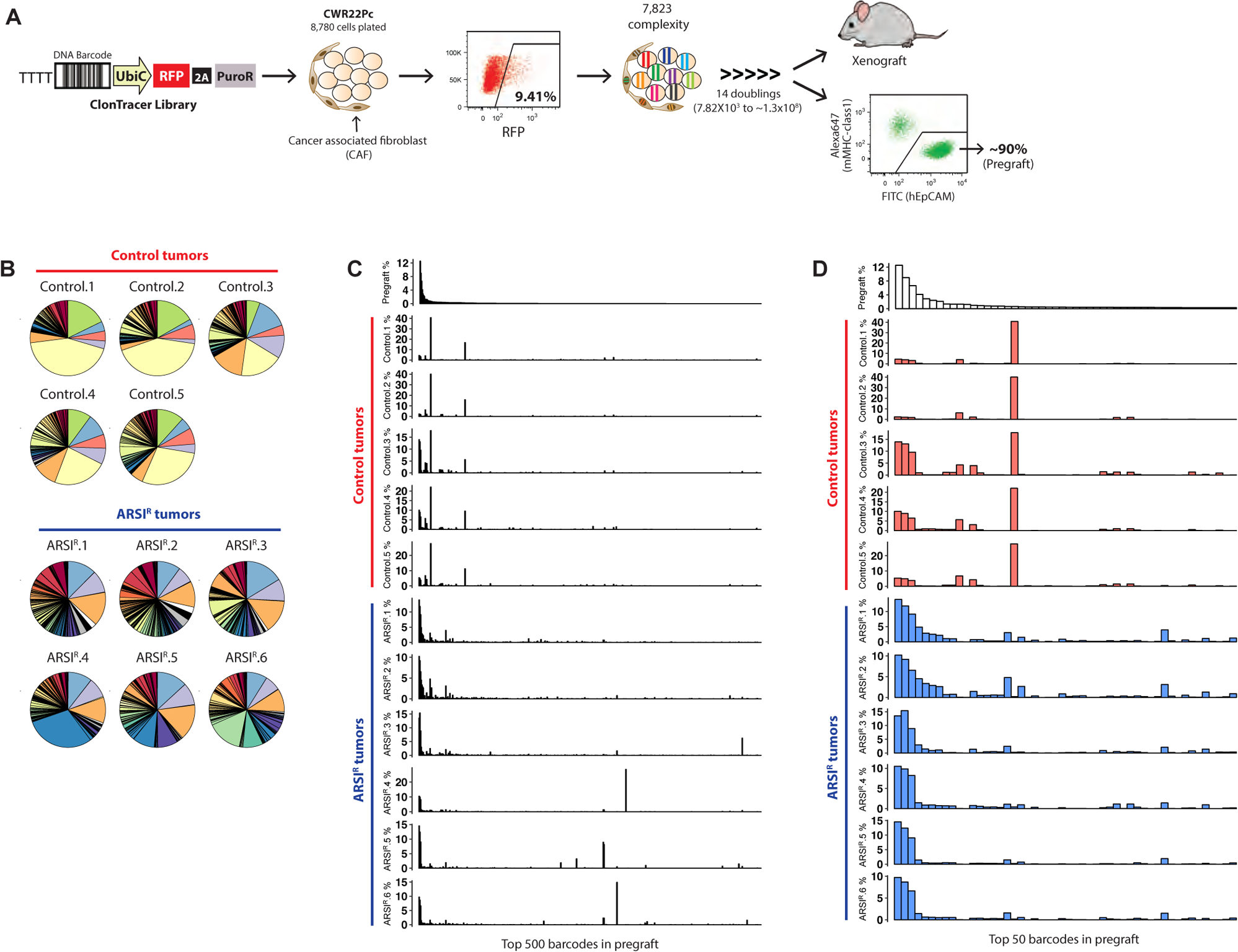
Enrichment of pre-existing persister clones in pre-engraftment ARSI setting in CWR22Pc. (A) The CWR22Pc cells containing mouse-derived cancer associated fibroblast (CAF) were infected with ClonTracer library to generate cell line labelled with 7,823 unique barcode, and then went through 14 doublings to have enough number for xenograft experiments. Since the CAFs are also labelled with barcode, we FACS sorted the human EpCAM-positive/mouse MHC-class1-negative population from the cells used for injection at each xenograft experiment and referred to it as ‘pregraft’. (B-D) Pie charts (B) and bar graphs (C-D) showing barcode distribution in control or ARSI-resistant (ARSI^R^) tumors from Fig. 2B. In bar graphs, the barcodes in x-axes are identical across all the graphs in a decreasing order of abundance of 500 (C) or 50 (D) most enriched barcodes in pregraft.

**Supplementary figure 5.**
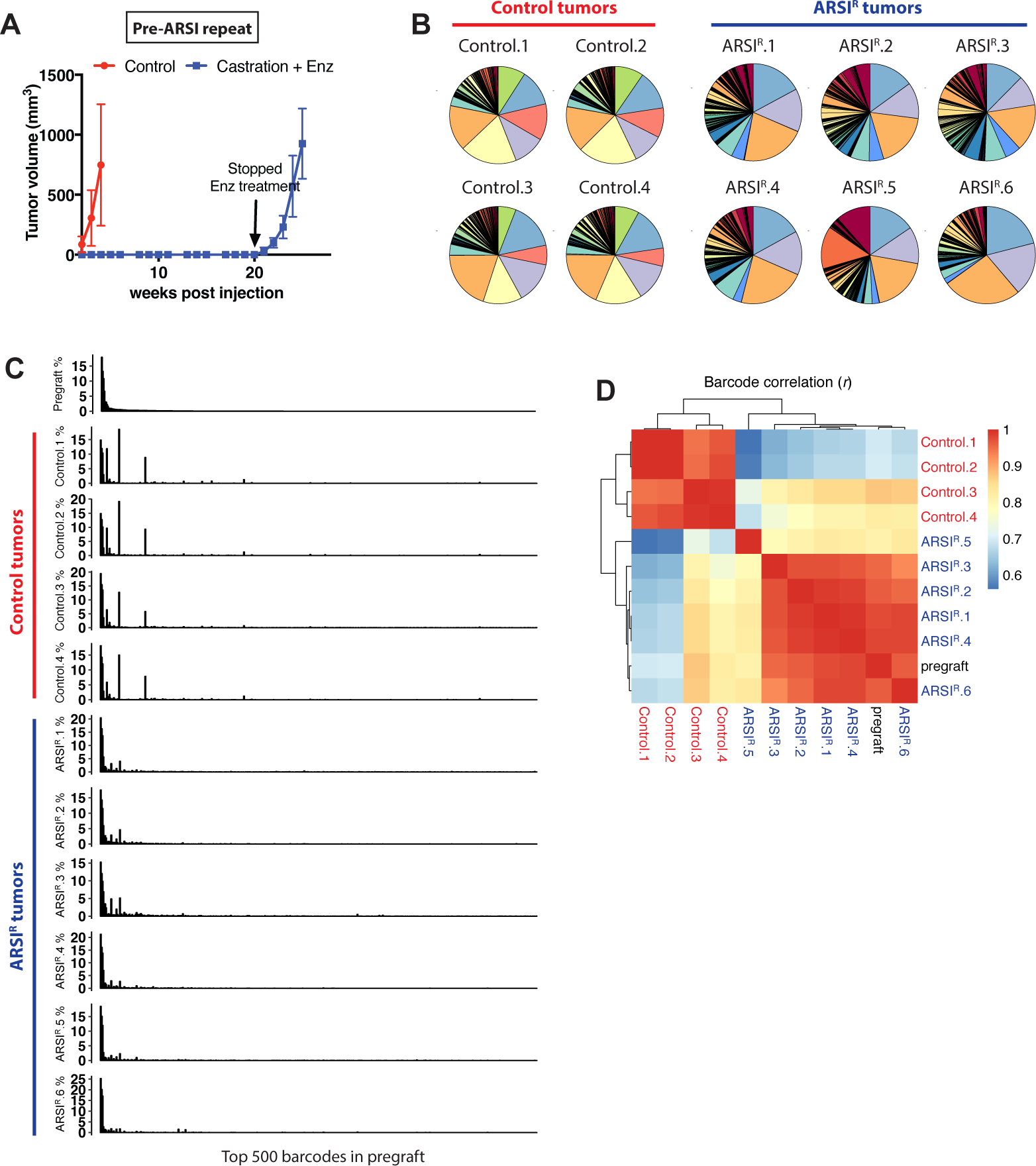
Replicate experiment showing enrichment of pre-existing resistance clones in setting of pre-engraftment ARSI therapy. (A) Mean tumor volumes ± SEM of the repeated pre-engraftment ARSI experiment (N=6 in control and N=11 in Enz treated group). Note that the Enz treatment is stopped at 20 weeks after grafting. (B-C) Pie charts (B) and bar graphs (C) showing barcode distribution in control and ARSI^R^ tumors. In bar graphs, the barcodes in x-axes are identical across all the graphs in a decreasing order of abundance of 500 most enriched barcodes in pregraft. (D) Heatmap depicting Pearson correlation analysis (*r*) of barcodes between the tumors shows that the control and ARSI^R^ tumors are clustered separately.

**Supplementary figure 6.**
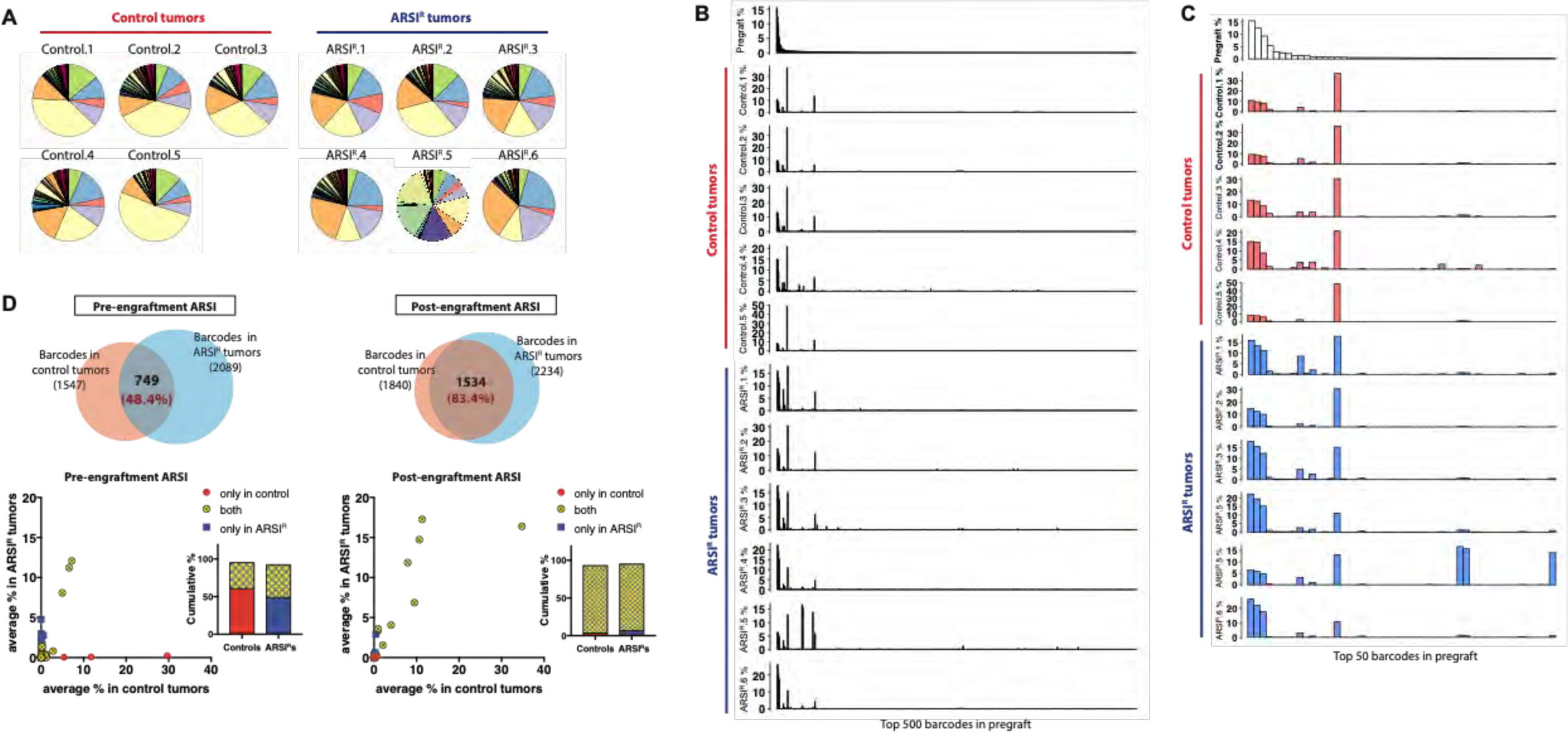
ARSI resistance in established CWR22Pc tumors is adaptive despite the presence of pre-existing resistant clones. (A-C) Pie charts (A) and bar graphs (B-C) showing barcode distribution in control and ARSI^R^ tumors from Fig. 2G. In bar graphs, the barcodes in x-axes are identical across all the graphs in a decreasing order of abundance of 500 (B) or 50 (C) most enriched barcodes in pregraft. (D) Top, Venn diagrams showing the overlaps between barcodes enriched in control and ARSI^R^ tumors in pre-engraftment and post-engraftment ARSI studies in Fig. 2. The % overlap is based on the number of barcodes enriched in control tumors. Bottom, scatter plots showing the average % of each barcode in tumors and bar graphs showing cumulative % of barcodes enriched only in control (blue), ARSI^R^ (red) or both groups (yellow).

**Supplementary figure 7.**
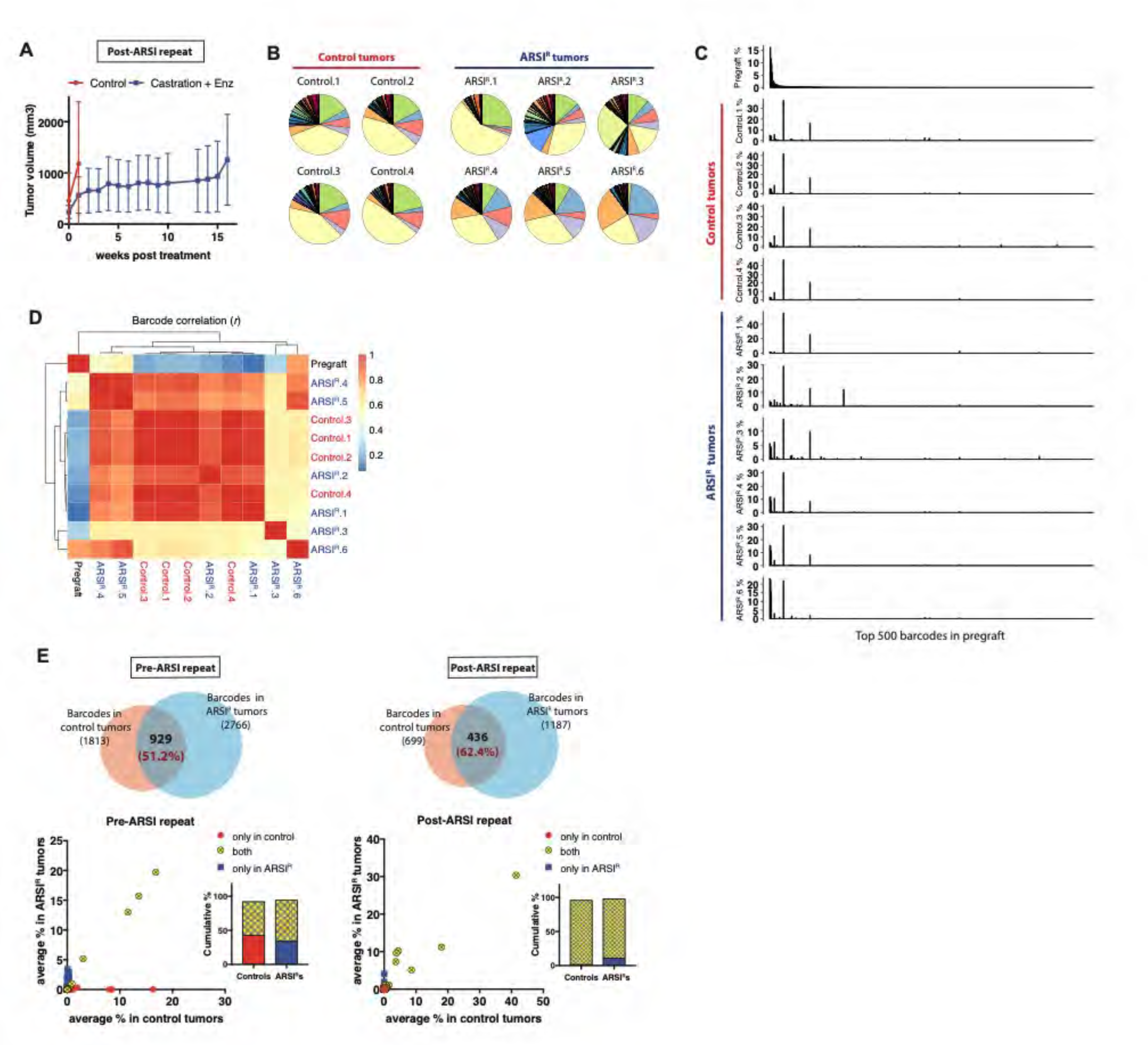
Replicate experiment confirming that ARSI resistance in established CWR22Pc tumors is adaptive despite the presence of pre-existing resistant clones. (A) Mean tumor volumes ± SEM of the repeated post-engraftment ARSI study (N=8 in control and N=6 in Enz treated group). (B-C) Pie charts (B) and bar graphs (C) showing barcode distribution in control and ARSI^R^ tumors. In bar graphs, the barcodes in x-axes are identical across all the graphs in a decreasing order of abundance of 500 most enriched barcodes in pregraft. (D) Pearson correlation analysis (*r*) of barcodes between the tumors shows that the control and ARSI^R^ tumors are clustered together. (E) Top, Venn diagrams showing the overlaps between barcodes enriched in control and ARSI^R^ tumors in repeated pre-engraftment and post-engraftment ARSI studies (Supplementary Figs. 5 and 7). The % overlap is based on the number of barcodes enriched in control tumors. Bottom, scatter plots showing the average % of each barcode in tumors and bar graphs showing cumulative % of barcodes enriched only in control (blue), ARSI^R^ (red) or both groups (yellow).

**Supplementary figure 8.**
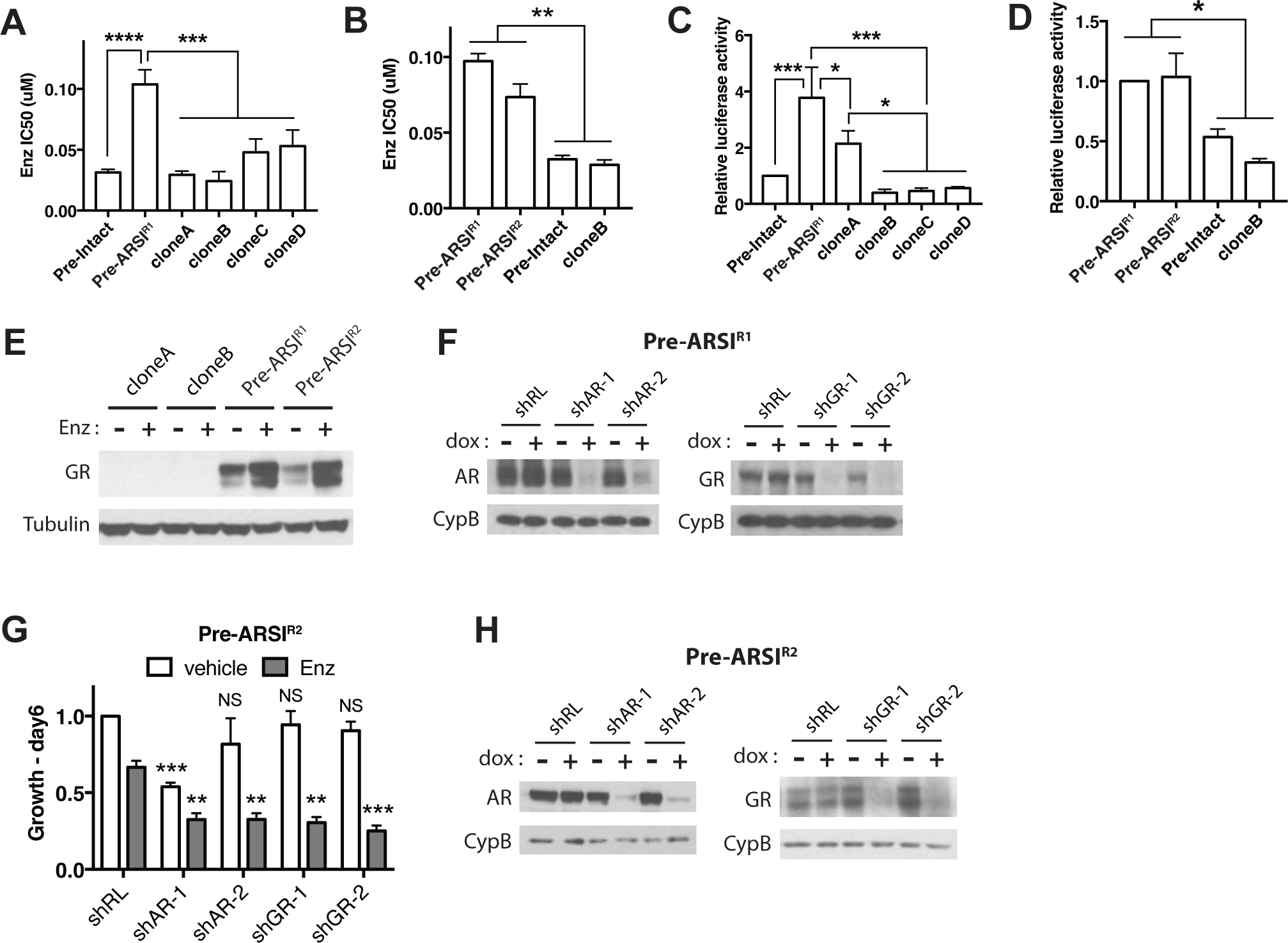
Pre-existing ARSI-resistant LNCaP/AR clones have higher baseline AR activity and GR expression. (A-B) Pre-ARSI^R1^ and Pre-ARSI^R2^ clones have higher IC50 values of Enz compared to other clones. IC50 values ± SD of three biological replicates. *P*-values were determined by one-way ANOVA. (C-D) Pre-ARSI^R1^ and Pre-ARSI^R2^ clones have higher AR reporter activities compared to the other clones. The values of luciferase AR reporter activities are normalized to Pre-Intact clone ± SD of three biological replicates. *P*-values were determined by one-way ANOVA. (E) GR is upregulated in both Pre-ARSI^R1^ and Pre-ARSI^R2^ clones and Enz treatment further increase GR expression. (F) The knockdown level of AR and GR from the cells in Fig. 3E. (G) Knockdown of AR or GR re-sensitize Pre-ARSI^R2^ clone to 10μM Enz. The cell viability values were normalized to the value of vehicle treated shRenilla (shRL) ± SD of two biological replicates. *P*-values were determined by two-way ANOVA compared to shRL. (H) The knockdown level of AR and GR from the cells in G. For all panels, NS = not significant, \**p*<0.05, \*\**p*<0.01, \*\*\**p*<0.001, \*\*\*\**p*<0.0001.

**Supplementary figure 9.**
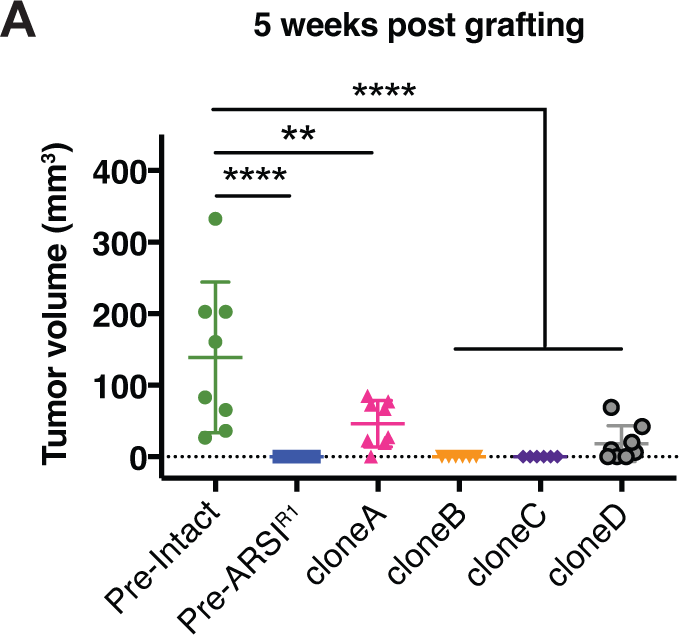
Mean tumor volumes ± SEM (N=8 in each group) at 5 weeks after grafting the 6 LNCaP/AR clones into hormonally intact mice. \*\**p*<0.01, \*\*\*\**p*<0.0001, one-way ANOVA.

